# Electrical Impedance Spectroscopy Quantifies Skin Barrier Function in Organotypic In Vitro Epidermis Models

**DOI:** 10.1101/2024.03.18.585587

**Authors:** N.J.M van den Brink, F. Pardow, L.D. Meesters, I. van Vlijmen-Willems, D. Rodijk-Olthuis, H. Niehues, P.A.M. Jansen, S. H. Roelofs, M.G. Brewer, E.H. van den Bogaard, J.P.H. Smits

## Abstract

3 D human epidermal equivalents (HEEs) are a state-of-the-art organotypic culture model in pre– clinical investigative dermatology and regulatory toxicology. Here, we investigated the utility of electrical impedance spectroscopy (EIS) for non–invasive measurement of HEE epidermal barrier function. Our setup comprised a custom–made lid fit with 12 electrode pairs aligned on the standard 24–transwell cell culture system. Serial EIS measurements for seven consecutive days did not impact epidermal morphology and readouts showed comparable trends to HEEs measured only once. We determined two frequency ranges in the resulting impedance spectra: a lower frequency range termed EIS^diff^ correlated with keratinocyte terminal differentiation independent of epidermal thickness and a higher frequency range termed EIS^SC^ correlated with *stratum corneum* thickness. HEEs generated from CRISPR/Cas9 engineered keratinocytes that lack key differentiation genes *FLG*, *TFAP2A, AHR* or *CLDN1* confirmed that keratinocyte terminal differentiation is the major parameter defining EIS^diff^. Exposure to pro–inflammatory psoriasis– or atopic dermatitis–associated cytokine cocktails lowered the expression of keratinocyte differentiation markers and reduced EIS^diff^. This cytokine–associated decrease in EIS^diff^ was normalized after stimulation with therapeutic molecules. In conclusion, EIS provides a non– invasive system to consecutively and quantitatively assess HEE barrier function and to sensitively and objectively measure barrier development, defects and repair.

## INTRODUCTION

Intact physical barriers are of highest importance for our body to define a biophysically enclosed environment. The skin, our largest barrier organ, serves a dual role: it forms an outside–in barrier, protecting the insides of our body from mechanical damage and environmental triggers, and it protects the epidermis and subjacent tissues from dehydration as an inside–out barrier. Barrier functionality is achieved most prominently by a highly organized physical barrier, constituted of tight junctions in the *stratum granulosum* and corneodesmosomes in the *stratum corneum* (Natsuga 2014). *Stratum corneum* corneocytes also are coated with a heavily crosslinked cornified envelope (Évora *et al*. 2021) and the intercellular space between is filled with lipids, generating a hydrophobic environment (van Smeden *et al*. 2014). Next to this physical barrier, the additional chemical, microbial and immunological barriers completes the multifaceted barrier function of mammalian skin (Eyerich *et al*. 2018; Niehues *et al*. 2018).

The importance of the skin barrier is apparent from its malfunction in common skin diseases, like psoriasis and atopic dermatitis. The disease–associated pro–inflammatory milieu also negatively affects keratinocyte differentiation and impairs tight junction and corneodesmosome function (Al Kindi *et al*. 2021; Orsmond *et al*. 2021; Yoshida *et al*. 2022). Next to these multifactorial diseases, monogenic diseases caused by mutations in skin barrier–related genes illustrate the devastating effects of impaired skin barrier function on our health and wellbeing (Hadj-Rabia *et al*., 2004; Supplemental Table 1 in van den Bogaard *et al*., 2019). Aside from these intrinsic factors, environmental factors including exhaust fumes or detergents influence the skin barrier function of healthy individuals and patients (Celebi Sözener *et al*. 2020). Determining the functional consequences of such genetic and environmental risk factors on the skin barrier will aid in our understanding of disease pathogenesis and may help in the possible future prevention of disease onset or exacerbation.

To investigate skin barrier function, *in vitro* organotypic skin and human epidermal equivalents (HEEs) have become a mainstay approach. By mimicking epidermal barrier morphology and function, HEEs offer advantages over *in vitro* monolayer cultures that lack epidermal stratification and *stratum corneum* formation. In addition, HEEs are considered relevant alternatives to *in vivo* animal models that prompt ethical questions and require depilation to measure biophysical barrier function. HEEs are used from fundamental research to preclinical drug testing to regulatory toxicology in a broad range of applications (El Ghalbzouri *et al*. 2008; Niehues *et al*. 2018).

To assess skin barrier function in HEEs, various technologies can be used ranging from mathematical penetration modelling (Pecoraro *et al*. 2019; Roberts *et al*. 2021) and computational simulation of lipid organization (Shamaprasad *et al*. 2022) to ultrastructural imaging (Riethmüller 2018) and measuring gene and/or protein expression. In a recent consensus paper, we and others have discussed the requirements and methodologies for barrier studies in organotypic skin models (van den Bogaard *et al*., 2021). In summary, functional barrier assessment using Franz cell diffusion and permeation flux studies provide most accurate estimates of the outside–in barrier (Arlk *et al*. 2018; Neupane *et al*. 2020; Tárnoki-Zách *et al*. 2021). On the other hand, water evaporation (e.g. trans–epidermal water loss (TEWL)) is considered most relevant to describe inside–out barrier function (Alexander *et al*., 2018; van den Bogaard *et al*., 2021). Unfortunately, these methods are often labor intensive and rely on highly specific expertise and equipment (Franz cell diffusion assay), are poorly standardized (transepithelial electrical resistance (TEER), TEWL) or require destructive endpoint measurements (permeation studies) (Table S1). Furthermore, the mechanistic correlation of such measurements to skin barrier function often remains unclear (Yoshida *et al*. 2022).

Electrical impedance spectroscopy (EIS) has been developed and implemented for the assessment of skin barrier function *in vivo* and appears to correlate well with disease severity of atopic dermatitis lesions (Rinaldi *et al*. 2021). For assessment of *in vitro* barrier function, EIS has been implemented for gut, airway and neuroepithelial *in vitro* cultures (Fernandes *et al*. 2022; Wegener *et al*. 2000) and *ex vivo* pig ear skin models (Morin *et al*. 2020). Explorative studies have applied EIS in *in vitro* epidermis models (Groeber *et al*. 2015; Mannweiler *et al*. 2021) and link EIS to viable epidermis and *stratum corneum* barrier properties (Mannweiler et al. 2021). Yet, a comprehensive study which extensively assesses EIS applicability and its relation to skin barrier properties in a broad range of experimental models and disease conditions is missing. Here, we demonstrate and validate the use of EIS as a reproducible, non–invasive and quantitative high– throughput system for HEE barrier assessment and assess its correlation to epidermal barrier physiology.

## RESULTS

### Development of an EIS device for *in vitro* HEE application

For quantitative and reproducible *in vitro* skin barrier analysis we sought to develop and validate an EIS devise for use in HEEs. The system comprises a smart lid with fixed gold–plated electrodes that are customized to fit the individual wells of a Nunc carrier plate with cell culture inserts. The setup enables standardized and automated measurements with a run time of 2 min per well for a maximum of 12 wells (within a 24 well–plate format). To perform the measurements, the smart lid with fixed electrodes is placed onto the HEE carrying culture plate and the connected measurement device (Figure 1a). The electrodes apply a very low alternating voltage *V* in a frequency range from 10 Hz to 100 kHz through the culture while measuring the amplitude and phase shift of the resulting alternating current *I*. The EIS device returns the impedance *Z*, which reflects the opposition to an alternating current over time:

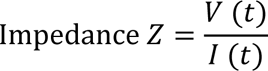

**Figure 1:**
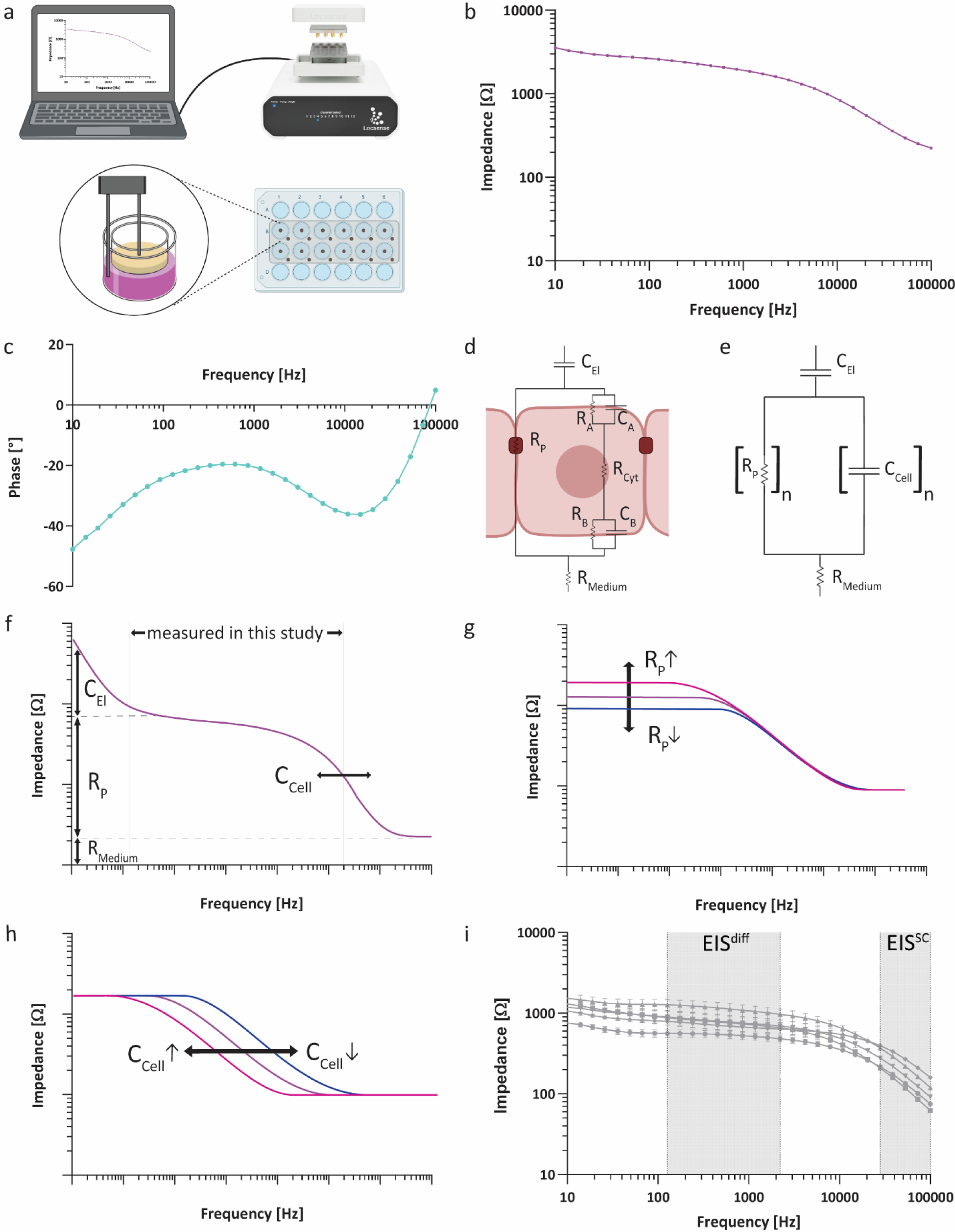
Design and function principle of a custom–made EIS device fitting the HEE culture system. (a) Schematic overview of the EIS setup on HEE cultures. (b) Impedance and (c) phase spectrum of a fully–developed HEE culture after 8 days of air exposure. (d) Extended electrical equivalent circuit of an epidermal monolayer culture made up of the capacitance of the electrodes C_El_, paracellular resistance R_P_, transcellular resistance of cytoplasm R_Cyt_, apical and basolateral membrane R_A_, R_B_ as well as their capacitance C_A_, C_B_ and the resistance of the medium R_Medium_ (adapted from (Yeste et al. 2018)). (e) Simplified electrical equivalent circuit of a HEE with the capacitance of both membranes taken together as C_Cell_ which together with R_P_ extends to a series of n parallel circuits in multi–layered 3 D culture systems (adapted from (Srinivasan et al. 2015) and (Groeber et al. 2015)). (f) Schematic overview indicating the contribution of individual electrical circuit parameters to the impedance spectrum (adapted from (Benson et al. 2013)). (g, h) Simulated impedance spectra illustrating the influence of changes in (g) paracellular flux (R_P_) and (h) transcellular flux (C_Cell_) (adapted from (Yeste et al. 2018)). (i) EIS impedance spectrum displaying EIS^diff^ (127**–**2212 Hz) and EIS^SC^ (28072**–**100000 Hz).

Impedance and phase spectra are reported in the form of a Bode plot (Figure 1b, c).

For quantitative analysis of EIS spectra, an electrical equivalent circuit model of the examined culture system is required. In conventional 2 D monolayer cultures, there are two main routes of the current: a paracellular, which is determined by the ionic conductance of cell junctions serving as a resistance R_P_, and a transcellular, which consists of the resistance and capacitance of apical and basolateral membrane (R_A_, R_B_ and C_A_, C_B_) next to the resistance of the cytoplasm R_Cyt_ (Benson *et al*. 2013; Srinivasan *et al*. 2015) (Figure 1d). In a simplified model, both membranes can be reduced to R_Cell_ and C_Cell_ respectively. In 2 D monolayers, the high cellular resistance R_Cell_ and the low cytoplasmatic resistance R_Cyt_ results in paracellular flux being determined by the cellular capacitance C_Cell_ (Benson *et al*. 2013). In 3 D HEEs, multiple individual cell layers result in a parallel series of n resistor–capacitor electrical circuits (Groeber *et al*. 2015). While we speculate R_P_, R_Cyt_, R_Cell_ and C_Cell_ to be changing depending on cell–cell contacts, differentiation status and the cell shape in the corresponding layer, we assume the dominance of C_Cell_ over R_Cell_ and R_Cyt_ to be persistent in 3 D (Figure 1e). Lastly, the resistance of the medium R_Medium_ and the electrodes, acting as pure capacitors with a capacitance C_El_, conclude the electrical circuit.

These electrical circuit elements also determine the generated impedance spectrum (Figure 1f) (Benson *et al*. 2013). Both, R_Medium_ and C_El_ are fixed parameters whose characteristics are determined by the chosen setup and device. The variable parameters paracellular resistance R_P_ and cellular capacitance C_Cell_ are determined by the measured cells and the chosen culture system (monolayer vs 3 D organoid). They influence the height and frequency span of a mid–range plateau and the onset of its decline (Yeste *et al*. 2018) (Figure 1g, h). To link EIS to epidermal barrier properties, we determined two frequency ranges in the HEE impedance spectrum approximately indicating R_P_ and C_Cell_ contribution, EIS^diff^ (127**–**2212 Hz) and EIS^SC^ (28072**–**100000 Hz) respectively, which we analyzed by calculating their area under the curve (Benson *et al*. 2013) (Figure 1i).

### Serial measurements show increased electrical resistance without affecting HEE **development**

To determine whether EIS can be used to monitor the development of HEEs non–invasively, serial measurements were performed over consecutive days of immortalized N/TERT–2G keratinocyte air–liquid interface culture. When comparing serial with end point measurements performed directly before harvesting, EIS spectra showed similar trends (Figure 2a, b) and differences in EIS^diff^ and EIS^SC^ were not significant (Figure 2c, d). Morphological analysis by hematoxylin and eosin (H&E) staining did not indicate destructive effects of EIS in endpoint or in serial measurements (Figure 2e, top panel). In addition, neither keratinocytes proliferation capacity (Ki67 staining), expression of differentiation proteins (involucrin (IVL) and filaggrin (FLG)), nor the expression of stress–related markers (keratin 16 (KRT16) and skin–derived antileukoprotease (SKALP)) were changed by EIS measurements (Figure 2e). Of note, expression levels of SKALP and the proliferation marker KRT16 are known to be higher in neonatal–derived immortal N/TERT–2G than in primary adult keratinocytes (Smits et al. 2017; Tjabringa et al. 2008). To further evaluate EIS’ reliability, HEEs were subjected to EIS measurements six times within one hour (Figure 2f), clearly indicating a very high repeatability. When checking for potentially delayed cytotoxic effects, HEEs harvested 24 hours after repeated EIS measurements showed no morphological signs of cytotoxicity and continuing maturation, as seen by the formation of additional epidermal layers (Figure 2g).

**Figure 2:**
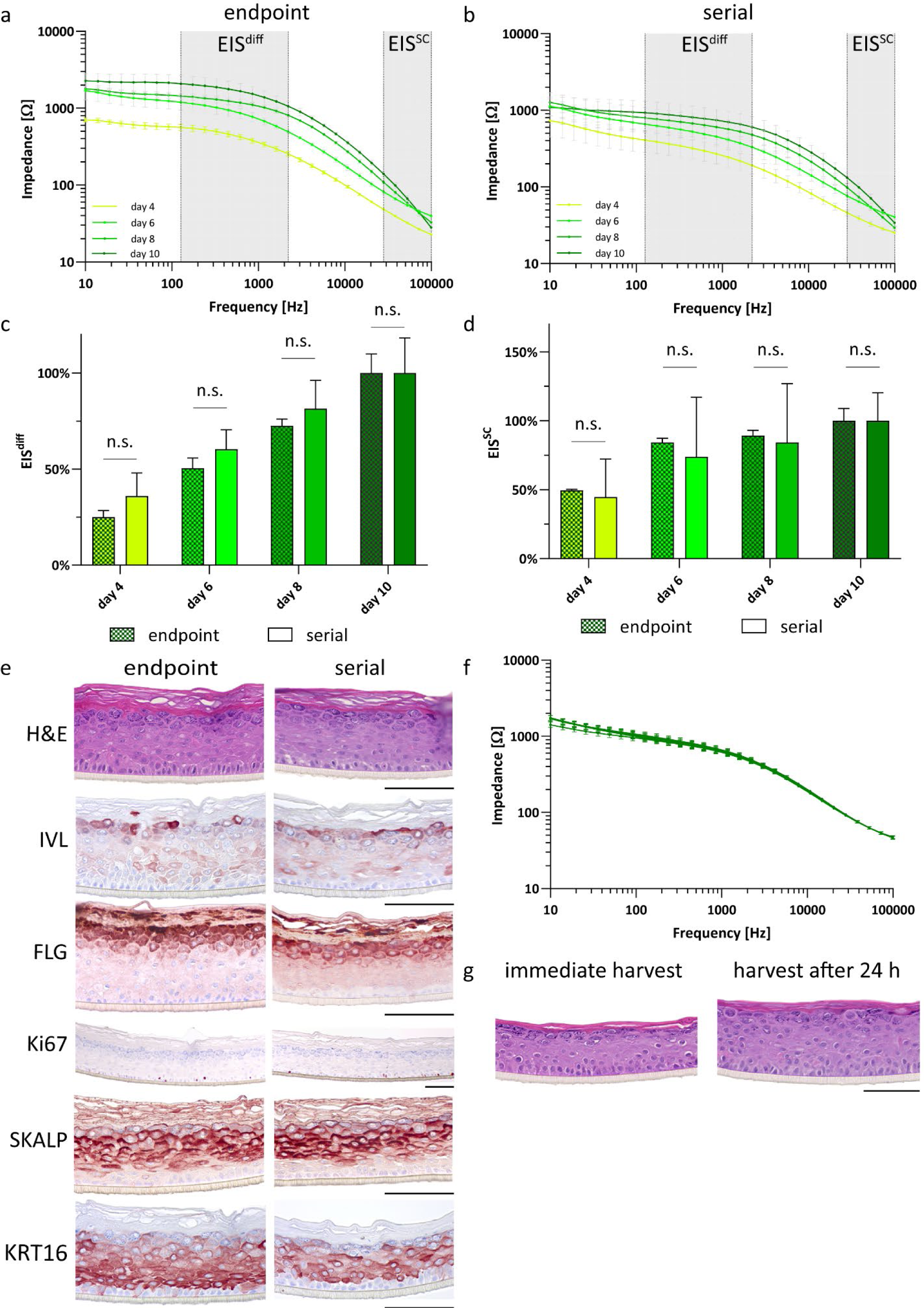
Relative EIS measurements are reproducible and do not impair HEE development. (a, b) EIS impedance spectra during HEE development with constructs being harvested (a) directly after measurements (endpoint measurements) and (b) at day 10 of air exposure (serial measurements). Each timepoint averages three biological replicates. (c) Comparison of EIS^diff^ and (d) EIS^SC^ between endpoint and serial measurements. (e) Histological comparison of HEEs undergoing endpoint or serial EIS measurements based on general morphology (H&E staining), differentiation status (FLG, IVL expression), proliferation (Ki67) and stress response (SKALP, KRT16). Pictures represent three biological replicates at day 10 of air exposure and taken at either 20x (Ki67) or 40x magnification. Size bars indicate 100 µm. (f) Impedance spectrum of HEEs (n = 3) measured 6 times within 1 hour at day 6 of air exposure. (g) Histological comparison of HEEs measured 6 times in 1 hour, either harvested directly or 24 hours after EIS measurements. Pictures represent three biological replicates and were taken at 40x magnification. Size bar indicates 100 µm.

### Different electrical impedance spectra can be linked to keratinocyte differentiation and ***stratum corneum* thickness**

The EIS device measures impedance over a broad range of frequencies which we sought to correlate with epidermal barrier properties. For this, we determined EIS^diff^ and EIS^SC^ of HEEs during barrier development (Figure 3a–c) and performed correlation analysis with principle epidermal barrier compartments (Figure 3d**–**g). We observed the thickness of viable epidermal layers to be increasing from day 1–10 before decreasing at day 12 and 14 (Figure S1a, b), which correlated with EIS^diff^ but not with EIS^SC^ (Figure 3d, e). At the same time *stratum corneum* thickness did not correlate with EIS^diff^ but strongly correlated with EIS^SC^ explaining 62 % of its variance (Figure 3f–g). Since keratinocyte terminal differentiation plays a major role in the formation of the skin barrier, we also investigated the protein expression of essential terminal differentiation proteins in relation to EIS^diff^ (Figure 3jh–l). EIS^diff^ correlated with the quantified expression of keratinocyte differentiation markers FLG and IVL which increased in early days of HEE development before decreasing after maturation at day 14 (Figure 3 h–j). Expression of the tight junction proteins, claudin 1 (CLDN1) and claudin 4 (CLDN4) could not be linked to EIS^diff^ (Figure 3h, k–l). When investigating the contributions of FLG, IVL, CLDN1 and CLDN4 altogether, CLDN1 and CLDN4 were not observed contributing to EIS^diff^ during HEE development and did not add additional explanatory value to the correlation model (Figure S1c, d). The expression of FLG, IVL and their collaborative interaction on the other hand together could explain 76 % of EIS^diff^ (Figure S1e, f).

**Figure 3:**
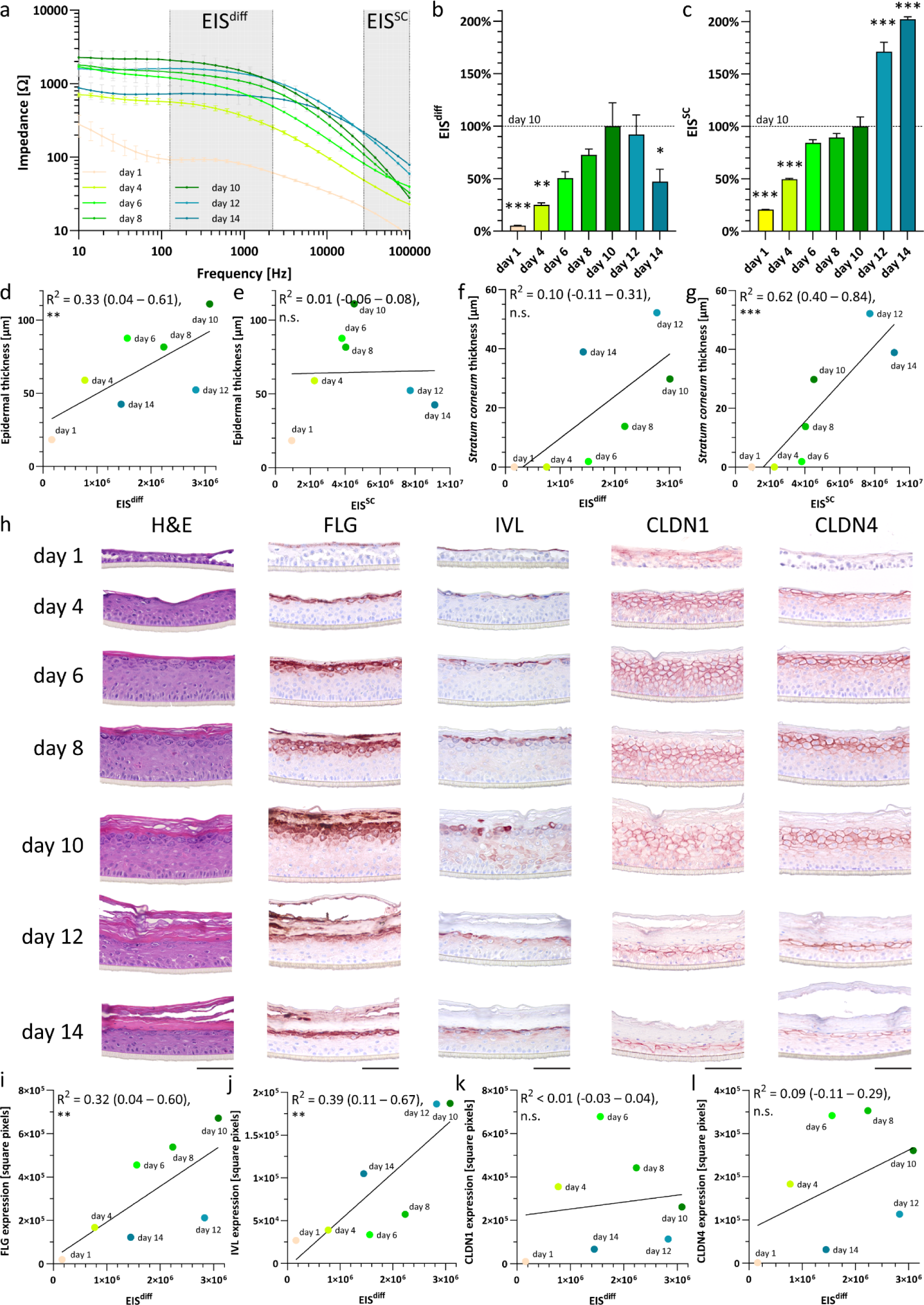
During HEE development, EIS^diff^ correlates with keratinocyte differentiation and epidermal thickness while EIS^SC^ correlates with *stratum corneum* thickness. (a) Endpoint– measured impedance spectra, (b) EIS^diff^ and (c) EIS^SC^ during HEE development. Each timepoint represents three biological replicates and EIS^diff^ and EIS^SC^ were compared to 10 day air–exposed cultures. (d – g) Correlation of epidermal thickness with (d) EIS^diff^ and (e) EIS^SC^ and *stratum corneum* thickness with (f) EIS^diff^ and (g) EIS^SC^. Each timepoint averages three biological replicates, R^2^ values and significances indicate the correlation of individual replicates. (h) Staining of general morphology (H&E), keratinocyte differentiation (FLG, IVL) and cell–cell adhesions (CLDN1, CLDN4) during HEE development. Pictures represent three biological replicates and were taken at 40x magnification. Size bars indicate 100 µm. (i – l) Correlation of (i) FLG, (j) IVL, (k) CLDN1 and (l) CLDN4 protein expression with EIS^diff^. Each timepoint averages three biological replicates, R^2^ values and significances indicate the correlation of individual replicates.

### Cytokine stimulation proves that EIS^diff^ is independent of epidermal thickness

After examining EIS in epidermal homeostasis we aimed to study the relevance of EIS in the context of disturbed homeostasis and to deepen our investigation into the correlation between EIS^diff^, epidermal thickness and terminal differentiation. Therefore, HEEs from normal human keratinocytes (NHEKs) were stimulated with single cytokines (interleukin–(IL–) 17A or IL–22) or cytokine mixes (IL–17A + IL–22 and IL–4 + IL–13) to mimic a pro–inflammatory milieu that is known to affect keratinocyte proliferation (IL–4, IL–13, IL–17A), the cell volume (IL–22), and terminal differentiation (all cytokines) (Niehues *et al*. 2021) (Figure 4a). We hypothesized that if EIS^diff^ would merely quantify epidermal thickness, cytokines known to increase epidermal thickness would increase EIS^diff^. Nevertheless, while IL–4 + IL–13 stimulation of HEEs significantly increased epidermal thickness, a reduction of EIS^diff^ was observed (Figure 4b, d).

**Figure 4:**
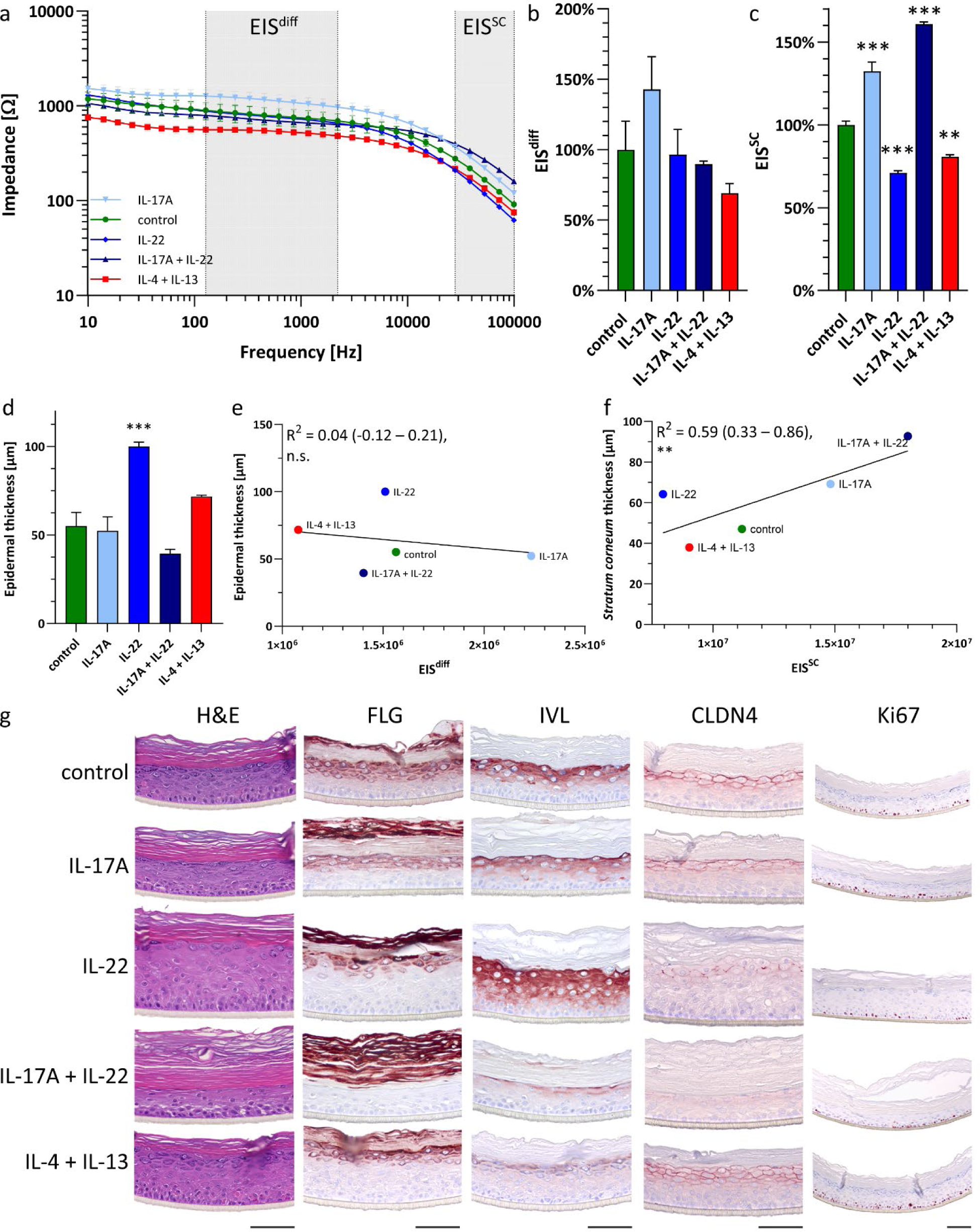
Stimulation with cytokines prove EIS^diff^–determined barrier function to be independent of epidermal thickness. (a) Endpoint–measured impedance spectra, (b) EIS^diff^ and (c) EIS^SC^ of cytokine–stimulated HEEs at day 8 of air exposure. Each condition represents three biological replicates and EIS^diff^ and EIS^SC^ were compared to control. (d) Epidermal thickness of cytokine–stimulated HEEs as compared to control. Correlation of (e) epidermal thickness to EIS^diff^ and (f) *stratum corneum* thickness to EIS^SC^. Each condition averages three biological replicates, R^2^ values and significances indicate the correlation of individual replicates. (g) HEEs stained for differentiation (FLG, IVL) and cell–cell adhesion (CLDN4) and proliferation (Ki67) proteins. Pictures represent three biological replicates and were taken at 20x (Ki67) or 40x magnification. Size bars indicate 100 µm.

Stimulation with IL–17A did not change epidermal thickness but resulted in increased EIS^diff^ and EIS^SC^ (Figure 4b**–**d). In contrast, IL–22 did not induce any changes in EIS^diff^, while significantly increasing the epidermal thickness (Figure 4b–d). Correlation analysis furthermore showed no correlation between epidermal thickness and EIS^diff^ (Figure 4e). To reassess the relationship between EIS^diff^ and terminal differentiation, we analyzed the expression levels of key terminal differentiation proteins FLG and IVL which are known to be reduced in human skin related to barrier defects, and known to be affected upon stimulation with IL–4 + IL–13 cytokines *in vitro* (Furue 2020). Indeed, decreased expression levels of FLG and IVL as well as unchanged levels of CLDN4 in HEEs treated with IL–4 + IL–13 cytokines (Figure 4g) corresponded to reduced EIS^diff^ (Figure 4b). On the other hand, IL–17A treatment, which is known to strengthen tight junction function (Brewer *et al*. 2019), significantly increased EIS^diff^, while epidermal thickness and FLG and IVL expression appeared unchanged (Figure 4b, d, g). This again indicates EIS^diff^ to quantify the complex dynamics of terminal differentiation and skin barrier formation rather than mirroring epidermal thickness. Quantifying FLG and IVL protein expression was not sufficient to model EIS^diff^ behavior (data not shown), indicating that the complex effect of cytokines on epidermal barrier function cannot be explained by the expression of FLG and IVL alone. EIS^SC^, however, was also found here to significantly correlate with *stratum corneum* thickness (Figure 4f).

### Knockout out of epidermal differentiation and cell–cell adhesion genes links EIS^diff^ to HEE differentiation

To further test our hypothesis that keratinocyte terminal differentiation significantly defines EIS^diff^, we knocked out key epidermal differentiation proteins by clustered regularly interspaced short palindromic repeats (CRISPR)/ CRISPR–associated protein 9 (Cas9)–mediated genome editing through non–homologous end–joining. We created keratinocyte cell lines lacking terminal differentiation protein *FLG* (Smits *et al*. 2023a), tight junction protein C*LDN1* (Arnold *et al*. 2023, accepted with minor revisions) or transcription factors known to coordinate terminal differentiation namely aryl hydrocarbon receptor (AHR) (Smits *et al*. 2023b, accepted) and transcription factor activating enhancer binding protein 2 alpha (TFAP2A) (Smits *et al*. 2023b, accepted). All knockout lines showed a reduction in EIS^diff^ and EIS^SC^ (Figure 5a–c). Notably, *CLDN1* knockout caused reduced EIS^diff^ but showed increased EIS^SC^ in concordance with observed parakeratosis, increased *stratum corneum* compaction (Figure 5d). FLG expression was clearly decreased in *AHR, TFAP2A* and *CLDN1* knockout lines and completely absent in the FLG knockout line (Figure 5d). Expression of IVL was decreased in all knockout cell lines, except the FLG knockout. CLDN1 (only fully absent in CLDN1 knockout) and CLDN4 expression appeared unchanged related to the epidermal cell layers which were affected by all genotypes as compared to control (Figure 5d). Hence, we conclude that EIS^diff^ is strongly determined by the degree of epidermal terminal differentiation.

**Figure 5:**
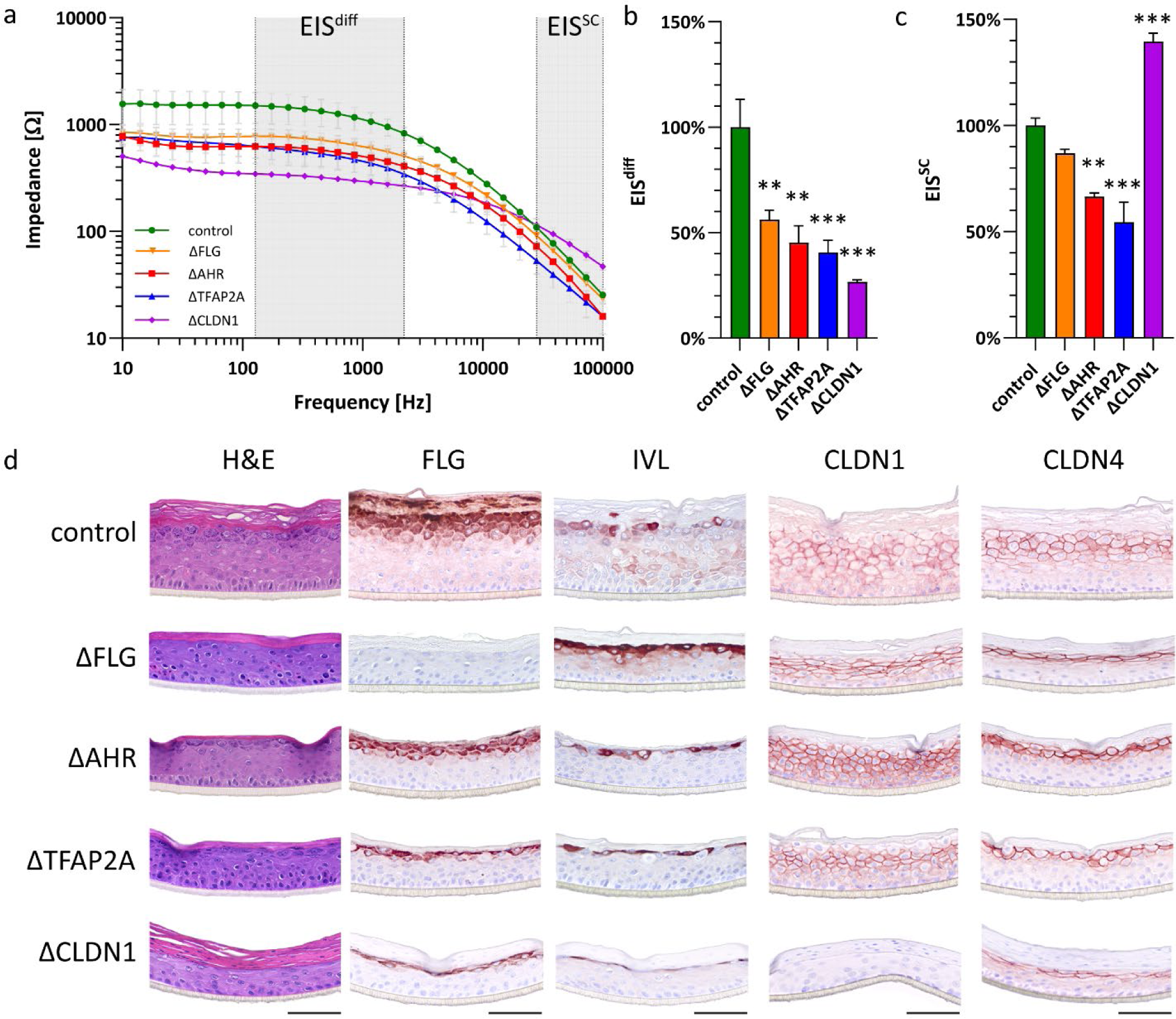
Knockout of genes involved in keratinocyte differentiation and cell–cell adhesion decreases EIS^diff^. (a) Endpoint–measured impedance spectrum, (b) EIS^diff^ and (c) EIS^SC^ of HEEs with knockout of target gene at day 10 of air exposure. Each condition represents three biological replicates and EIS^diff^ and EIS^SC^ are compared to control. Each (d) HEEs stained for differentiation (FLG, IVL) and cell–cell adhesion (CLDN1, CLDN4) proteins. Pictures represent three biological replicates and were taken at 40x magnification. Size bars indicate 100 µm.

### EIS detects therapeutic response in pro–inflammatory IL–4 + IL–13 epidermis model

Besides detecting or monitoring epidermal defects, reversing these defects is a key component in the treatment of inflammatory skin diseases and an important parameter in the development of potential novel therapeutics. Therefore, we investigated if EIS can measure the reversal of barrier defects for future implementation in pre–clinical drug screening. For this we chose pharmacological molecules known to activate AHR, a key regulator of epidermal differentiation (Figure 5d) and novel target for topical anti–inflammatory treatment (Bissonnette *et al*. 2021; Van Den Bogaard *et al*. 2013). Hereto IL–4 and IL–13 stimulated HEEs were additionally treated with an array of AHR–activating ligands (L1–5) known to have a therapeutic effect, next to structurally–related non–activating compounds (M1–2) (Table S2). AHR–activating compounds resulted in restored IL–4 + IL–13–impaired EIS^diff^ and EIS^SC^ impedance spectra, indicating capability of EIS to measure the AHR–dependent repair of skin barrier defects (Figure 6a–c). Compounds that do not activate AHR signaling (M1–2) did not restore the cytokine–mediated reduction in EIS (Figure 6a–c). In fact, these compounds known to block endogenous AHR signaling and further decreased EIS^SC^ significantly with similar trends in EIS^diff^.

**Figure 6:**
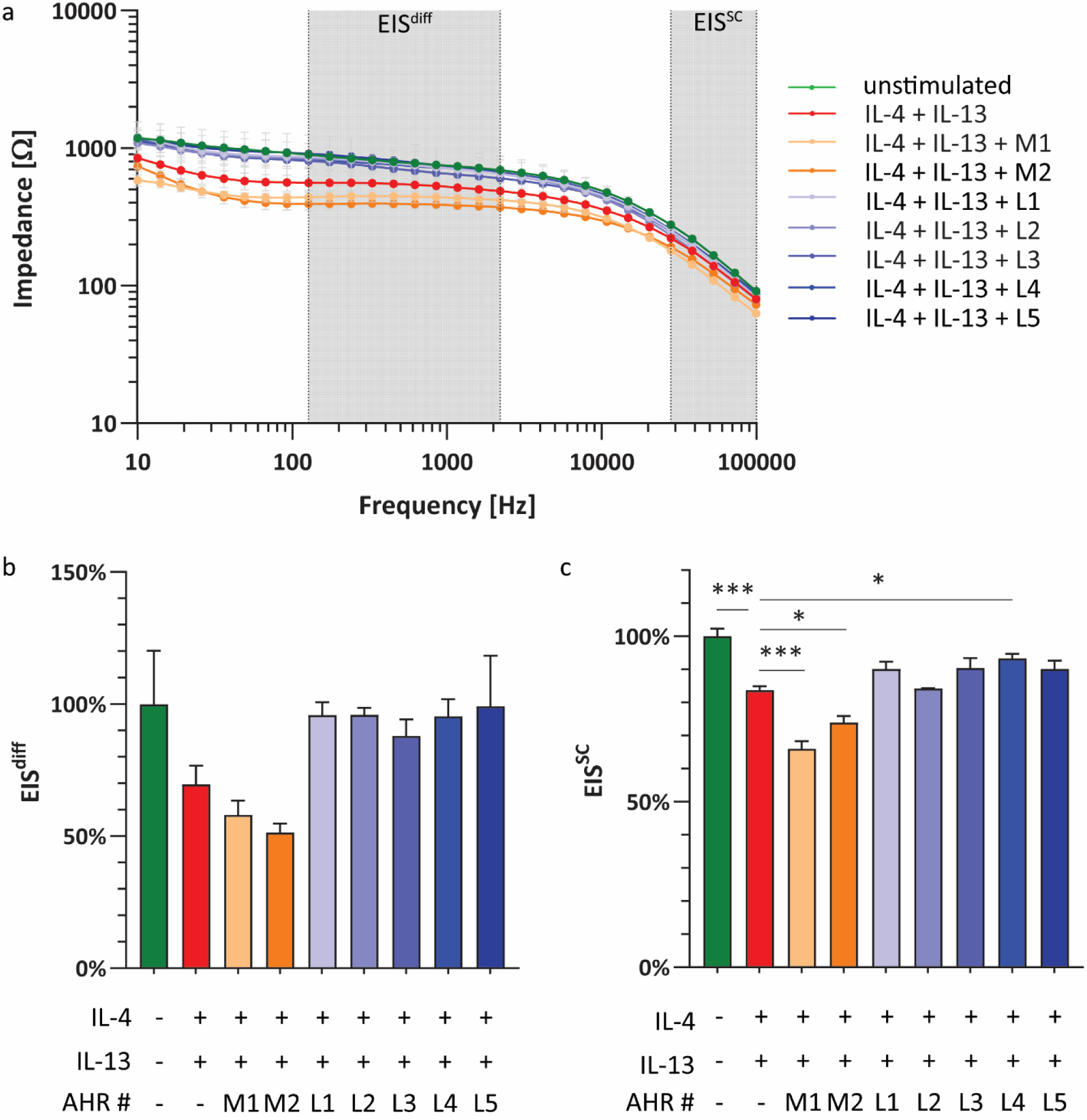
EIS detects therapeutic AHR response in a pro–inflammatory epidermis model. (a) Endpoint–measured impedance spectrum, (b) EIS^diff^ and (c) EIS^SC^ of HEEs stimulated with IL–4 and IL–13 cytokines alone and in combination with AHR activating therapeutic compounds at day 8 of air exposure. Each condition represents three biological replicates and EIS^diff^ and EIS^SC^ of control conditions and AHR–binding compounds are compared to IL–4 + IL–13 stimulation.

Since we now confidently determined that EIS^diff^ measurements correlate with keratinocyte differentiation, we assumed that expression levels of differentiation proteins FLG and IVL would also correlate with the rescue in EIS^diff^ by AHR ligands. This we could demonstrate clearly for IVL, as AHR agonists were partially able to restore the dampened IVL expression by IL–4 and IL–13. AHR–binding but non–activating compounds (M1–2) did not restore IVL expression (Figure S2a). FLG protein levels followed a similar expression pattern as IVL albeit differences were less pronounced. Again, the expression of tight junction protein CLDN4 remained unchanged throughout treatments (Supplemental Figure 2) confirming earlier results.

## DISCUSSION

In this study, we investigated the applicability of EIS to assess skin barrier function in 3 D HEE *in vitro* organotypic epidermis models. EIS proved an easy to handle and non–invasive system to obtain real–time quantitative readouts correlating to functional barrier properties.

While fit here to the Nunc carrier plate system, the device can be customized to various transwell cell culture platforms. Running costs are low and the measurements are performed in a semi– automated fashion. The fixed electrode setup additionally standardizes the measurements and produced results are highly repeatable when taking into account that electrical impedance readouts dependent on cell passage number, culture medium and temperature (Srinivasan *et al*. 2015). Reported readout trends are reproducible while absolute values have been observed to differ between replicated experiments. Sample readouts therefore need to be correlated with controls within the same experiment (see guideline box). While endpoint measurements cater to less variance, measurements can be performed and repeated in high quantities at virtually any time during HEE culture without harming tissue integrity. With current functional barrier assessments structurally relying on invasive endpoint measurements (permeation studies, Franz cell diffusion assay) and / or being laborious and sensitive to handling (Franz cell diffusion assay, TEWL, TEER), EIS provides a non–invasive, semi–automated and reproducible alternative.

EIS measures a broad range of frequencies in contrast to single frequency TEER measurements (Faway *et al*. 2021; Srinivasan *et al*. 2015) therefore being able to capture different functional skin barrier parameters. For quantitative analysis, experimentally obtained impedance spectra are usually fit to the corresponding electrical circuit model to isolate individual electrical parameters (Magar *et al*. 2021). To aid biologic interpretation, we here chose to correlate the obtained impedance spectra with known biological barrier properties. Theoretical considerations in combination with our own data highlighted two frequency ranges which we described through calculating their area under the curve.

Frequencies on a plateau around 100–2000 Hz were termed EIS^diff^ as they quantified differentiation in viable keratinocytes as assessed by the expression of differentiation markers FLG and IVL, which together predicted 76 % of EIS^diff^ during HEE development. While EIS^diff^ exhibits clear independence of viable epidermis and *stratum corneum* thickness, a correlation with tight junction function remains uncertain. On the one hand, EIS^diff^ was observed to be independent from CLDN1 and CLDN4 protein expression during HEE development and cytokine stimulations. ΔCLDN1 HEEs exhibits a strongly reduced EIS^diff^, but this can be explained by a concomitant reduced expression of keratinocyte differentiation markers as CLDN1 knockout is known to alter pro–FLG processing (Sugawara et al. 2013). On the other hand, EIS^diff^ frequencies overlap with R_P_ impedance contribution, with R_P_ being the electrical circuit model element to describe the resistance of tight junctions (Srinivasan *et al*. 2015). Furthermore, during IL–17A cytokine stimulation EIS^diff^ increased independent of differentiation protein expression, which could be a result of an IL–17A strengthening effect on tight junction function (Brewer *et al*. 2019). In conclusion, EIS^diff^ uniquely quantifies keratinocyte differentiation independent of CLDN1 and CLDN4 protein expression, however a contribution of tight junction function cannot be ruled out since sole protein expression does not entirely mirror tight junction functionality (Bäsler *et al*. 2016; Kirschner *et al*. 2012).

Frequencies of a higher frequency range between 20,000 Hz and 100,000 Hz were termed EIS^SC^ and overlap with C_Cell_, a parameter describing ability of a cell to store an electrical charge. EIS^SC^ conclusively quantifies *stratum corneum* more than complete HEE thickness and seems to be a descriptive rather than functional assessor since the thickened but parakeratotic ΔCLDN1 *stratum corneum* exhibited an increased EIS^SC^. We did not investigate EIS^SC^ dependence on lipid organization and *stratum corneum* composition, which should be the focus of future studies.

This study used EIS to assess HEE skin barrier function during formation, to study the effects of single genes and to assess skin barrier function under inflammatory conditions and treatment. EIS was able to measure the defects induced by knockout of cardinal differentiation–driving transcription factors (AHR and TFAP2A) and differentiation effector genes (FLG). The observed defects were congruent with other barrier function assessments reporting an elevated TEWL in ΔTFAP2A and ΔFLG HEEs and in human FLG loss–of–function variants (Nemoto-Hasebe *et al*. 2009; Smits *et al*. 2023b, accepted; Smits *et al*. 2023a; Yang *et al*. 2020). Cytokine–induced pro– inflammatory conditions resulted in keratinocyte differentiation deficiencies and changes in *stratum corneum* thickness which could be captured and quantified by EIS. The IL–4 + IL–13 induced decrease in EIS^diff^ and EIS^SC^ *in vitro* also replicates the *in vivo* situation where the IL–4 and IL–13 driven skin disease atopic dermatitis is accompanied by elevated TEWL and decreased EIS values measured on *in vivo* patient skin (Chamlin et al. 2002; Flohr et al. 2010; Rinaldi et al. 2021; Sugarman et al. 2003). *In vivo*, EIS can also detect therapeutic improvements of atopic dermatitis associated with improvements in clinical scoring and reduced expression of inflammatory biomarkers (Breternitz *et al*. 2008; Rinaldi *et al*. 2021) similar to the detected therapeutic improvements in our *in vitro* atopic dermatitis model.

To conclude, we propose EIS to be a valuable tool to non–invasively study epidermal barrier function in organotypic skin models. The dual viable epidermis/*stratum corneum* barrier assessment and the quantification of keratinocyte differentiation is to our knowledge singular across all barrier evaluation techniques. The proposed semi–quantitative EIS analysis is easy to replicate and uniquely correlates impedance readouts with biological barrier properties. We suggest EIS to be especially suited for longitudinal studies of barrier development, keratinocyte differentiation and barrier–disrupting skin diseases including pre–clinical therapeutic studies. In addition, EIS can be used in multi–cell type models to investigate the interplay between epidermis, extrinsic and intrinsic factors, potentially in combination with patient–derived cells, immune cells and/or bacteria. The possibility to correlate *in vitro* and *in vivo* EIS measurements facilitates a unique translational approach from bedside to bench and back.

## TECHNICAL RECOMMENDATIONS

To ensure optimal, reproducible EIS measurements without compromising culture integrity, we propose several guidelines for implementing EIS in the laboratory:

- Measurements depend on temperature and ion content of the surrounding fluid. To ensure maximum comparability between conditions, use an isotonic buffer solution (e.g. phosphate–buffered saline (PBS)) and allow samples to adjust to room temperature for at least 30 min before measurements.
- To minimize variation when performing serial measurements at various days of the cell culture, the time of topical exposure to PBS should be kept minimal and PBS should be carefully removed after measurements to maintain the air–liquid interface as much as possible for proper barrier formation and function.
- Before commencing measurements, blank measurements on PBS only or empty filters should be performed, as this provides information on intrinsic capacitance of the electrodes and the resistance of PBS and filters. When analyzing the results, blanks should be subtracted from measured sample values.
- Previous publications have normalized EIS based on the surface area of the used cell culture system using various methods (Chen et al. 2015; Haorah et al. 2008; Liu et al. 2020). Considering the lack of consensus, we report uncorrected EIS values and the surface area of HEEs (0.47 cm^2^) to aid comparisons.
- Control conditions should be taken along for each individual experiment and measurement time point to interpret relative changes in preference to absolute values.
- EIS^diff^ (127**–**2212 Hz) and EIS^SC^ (28,072**–**100,000 Hz) are determined through calculating the area under the curve at respective frequencies.

## MATERIALS & METHODS

### Cell culture

Human primary keratinocytes were isolated from surplus human skin obtained through plastic surgery according to the principles and guidelines of the principles of Helsinki. From the skin, biopsies were taken and keratinocytes were isolated as described previously (Tjabringa *et al*. 2008). N/TERT–2G keratinocytes were a kind gift of James Rheinwald, Brigham’s Woman hospital (Dickson *et al*. 2000) and were cultured as monolayers in CnT–prime (CELLnTEC, Bern, Switzerland, CnT–PR) until confluent before use in HEE cultures (Smits *et al*. 2017). Knockout N/TERT–2G cell lines were generated through CRISPR/Cas9 and validated previously (FLG (Smits *et al*. 2023a), CLDN1 (Arnold *et al*. 2023, accepted with minor revisions), TFAP2A (Smits *et al*. 2023b, accepted), AHR (Smits *et al*. 2023b, accepted)).

### Generation of HEEs

HEE cultures were performed as previously described (Smits *et al*. 2023a). In short, cell culture inserts in a 24 wells carrier plate (Nunc, Thermo Fisher Scientific, 141002) were coated using 100 µg/mL rat tail collagen (Sigma–Aldrich, C3867) for 1 hour at 4 °C. After phosphate–buffered saline (PBS) washing the filters, 150000 cells were seeded and submerged in CnT–prime medium (CELLnTEC, CnT–PR) at the lowest insert stand. After 48 hours, the medium was switched to differentiation medium (40 % Dulbecco’s modified Eagle’s Medium (Sigma–Aldrich, D6546) and 60 % 3 D barrier medium (CELLnTEC, CnT–PR–3D)) and 24 hours afterwards the HEEs were lifted to the highest stand, air–exposed and medium was refreshed every other day. For stimulation experiments, IL–4, IL–13, IL–17A or IL–22 (50 ng/mL per cytokine, Peprotech, Rocky Hill, NJ, USA, 200-04/200-13/200-17/200-22) supplemented with 0.05 % bovine serum albumin (Sigma– Aldrich, A2153) were added to the medium of HEEs of primary keratinocyte from day 5 of air exposure until day 8. AHR ligands (Table S2) were supplemented in the culture medium as previously described (Rikken et al. 2023).

### EIS measurements

For the EIS measurements the Locsense Artemis (Locsense, Enschede, the Netherlands) device was used and equipped with a custom–made incubator compatible smart lid. The Artemis consists of a detector element that is connected to the smart lid with electrodes aligning to the two middle rows of a 24–well plate. A laptop equipped with the Locsense Artemis monitoring software (version 2.0) displays the readouts. During the measurements each well contains two electrodes: one disc–shaped 4.2 mm diameter electrode situated in the center of the transwell insert and a rod– shaped 1.9 mm diameter electrode passing sideways of the transwell insert. Before measurements, HEEs were acclimated to room temperature and cultures were lowered to the middle position in the transwell plate while 1600 μL PBS at room temperature was added below and 500 μL PBS on top of the filter. Thereafter, the smart lid was placed on the wells ensuring both electrodes being submerged. Following device self–calibration, impedance was measured over a frequency range from 10 Hz to 100,000 Hz in 30 logarithmic intervals. Measurement output contains impedance as well as phase values. Phase values can be interpreted as is while a PBS only blank measurement was subtracted from the corresponding electrode of the impedance output. For further specific considerations during measurements, see guideline box. For EIS^diff^ (127**–**2212 Hz) and EIS^SC^ (28,072**–**100,000 Hz) the area under the curve was calculated using the respective frequency ranges.

### Immunohistochemistry

For histological processing, 4 mm biopsies were fixated in 4 % formalin solution for 4 hours and embedded in paraffin. Afterwards, 6 µm sections were deparaffinized and either stained with hematoxylin (Klinipath, 4085.9005) and eosin (Klinipath, 4082.9002) or by antibodies listed in Table S3 followed by avidin–biotin complex (Vectastain, AK–5000). Epidermal thickness specifies the average of three measurements on H&E stained sample pictures while *stratum corneum* thickness was determined by subtracting epidermal from total construct thickness. Protein expression was quantified with ImageJ following sections C–E of (Crowe and Yue 2019) by freehand selecting the viable epidermis and measuring the “area” i.e. number of staining– positive square pixels.

### Statistics

Data sets were analyzed using the GraphPad Prism 10 software version 10.1.1. All barplots are shown as mean ± standard error of the mean and significance testing was performed using one– way analysis of variance (ANOVA) in combination with Dunnett correction for multiple testing and unpaired t–testing (exclusively in figure 2c, d). Differences under p value < 0.05 were considered statistically significant, ns p value > 0.05, * p value < 0.05, ** p value < 0.01, *** p value < 0.001.

### Correlation analysis of EIS to protein expression and HEE morphology

Correlation analysis was conducted using simple (epidermis thickness and *stratum corneum* thickness) and multiple linear regression modelling (protein expression) in Graphpad Prism and the R programming language (version 4.2.3) (R Core Team 2023) with the psychometric package (Fletcher 2023). All correlation analysis were conducted with individual replicates, figures depict replicate averages for readability.

## Abbreviations

ANOVA: analysis of variance
AHR: aryl hydrocarbon receptor
C_A_, C_B_: cellular membrane capacitance
C_Cell_: cellular capacitance
C_El_: electrode capacitance
CLDN1: claudin 1
CLDN4: claudin 4
CRISPR/Cas9: clustered regularly interspaced short palindromic repeats / CRISPR–associated protein 9
EIS: electrical impedance spectroscopy
EIS^diff^: keratinocyte differentiation–attributable electrical impedance
EIS^SC^: *stratum corneum*–attributable electrical impedance
H&E: hematoxylin and eosin
HEE: human epidermal equivalents
IL: interleukin
PBS: phosphate–buffered saline
R_A_, R_B_: cellular membrane resistance
R_Cell_: cellular resistance
R_Cyt_: cytoplasmatic resistance
R_Medium_: culture medium resistance
R_P_: paracellular resistance
TEER: transepithelial electrical resistance
TEWL: transepidermal water loss
TFAP2A: transcription factor activating enhancer binding protein 2 alpha
IVL: involucrin
FLG: filaggrin
KRT16: keratin 16
SKALP: skin–derived antileukoprotease

## DATA AVAILABILITY STATEMENT

Datasets related to this article are available from the corresponding author upon request.

## ACKNOWLEDGMENTS

We thank Joachim Wegener (University of Regensburg, Germany) for the critical reading of our manuscript and all members of the Van den Bogaard group for the lively discussions and suggestions. We thank Ewald Bronkhorst for advice on the correlation analysis. This collaborative work was supported by NIH R35 grant ES028244, PAST4FUTURE grant LSHM20043-HSGF and European Innovation Council (EIC) under grant agreement No. 101098826 (SKINDEV) and the Radboud university medical center (EB). The FLG knockout cells were generated under a LEO Foundation grant LF-OC-22-001056 (JS and EB).

## AUTHOR CONTRIBUTIONS

Conceptualization: NB, FP, LM, JS, and EB; Data Curation: NB, FP, and LM; Formal Analysis: NB and FP; Funding Acquisition: JS, EB; Investigation: NB, FP, LM, HN, IVW, MB, DRO, PJ JS; Methodology: NB, FP, LM, JS; Project Administration: EB; Software: SR; Supervision: JS and EB; Validation: NB and FP; Visualization: NB and FP; Writing – Original Draft Preparation: NB and FP; Writing – Review and Editing: all authors.

## SUPPLEMENTARY MATERIAL

**Supplemental Table 1.**
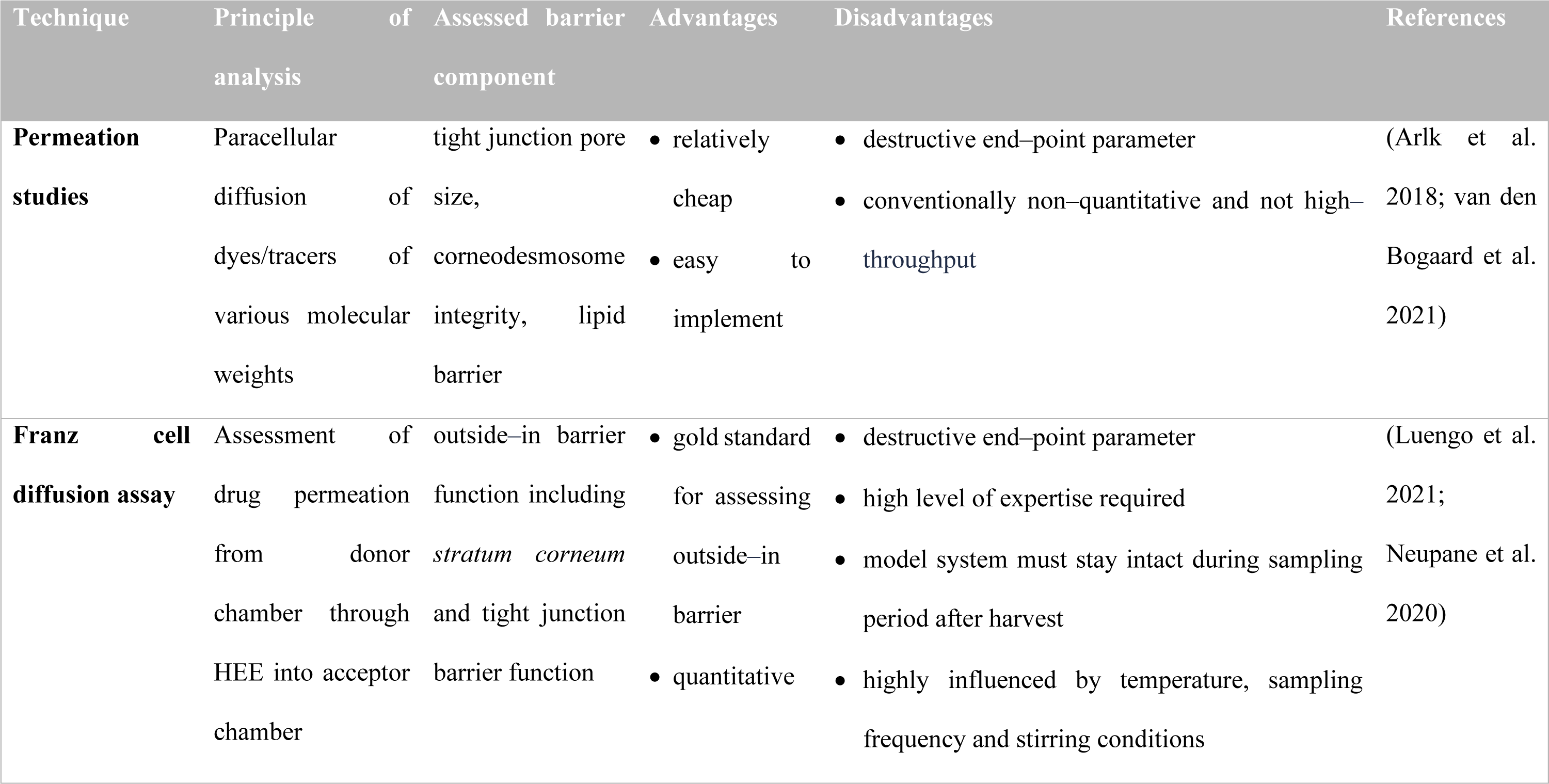
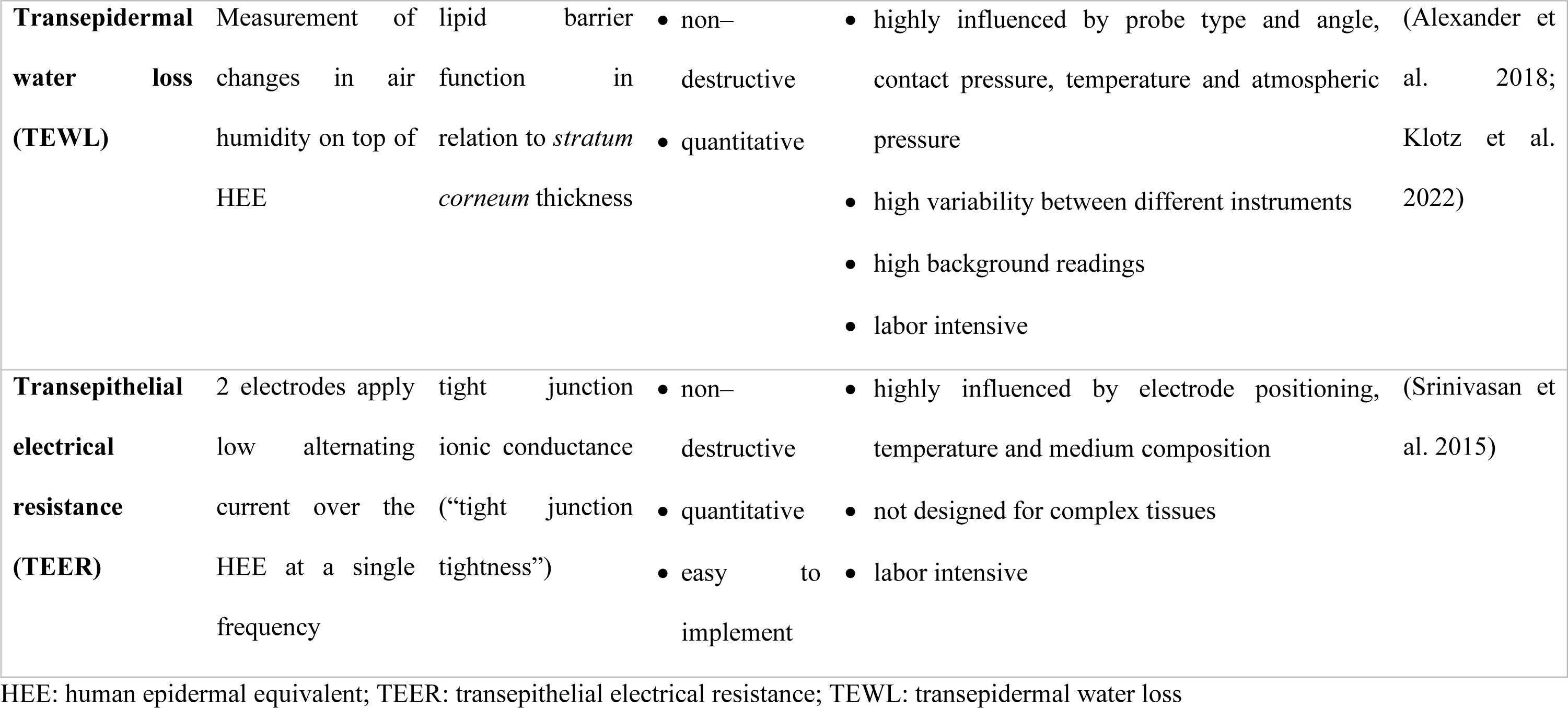
Overview of methodologies used for measurement of *in vitro* epidermal barrier function.

**Supplemental Table 2:** AHR–binding compounds.

| Abbreviation | Compound | Concentration | Reference |
| --- | --- | --- | --- |
| **M1** | SGA360 | 10 nM | (Rikken et al. 2023) |
| **M2** | Teriflunomide | 10 µM | (O'Donnell et al. 2010) |
| **L1** | coal tar | 2 µg/mL | (Van Den Bogaard et al., 2013) |
| **L2** | SGA388 | 10 nM | (Rikken et al. 2023) |
| **L3** | Leflunomide | 10 µM | (O'Donnell et al. 2010) |
| **L4** | IMA-7101 | 1 nM | (Rikken et al. 2022) |
| **L5** | TCDD | 10 nM | (Poland and Glover 1976; Safe 1986) |

**Supplemental Table 3:** Antibodies used for immunohistochemical analysis.

| Antigen | Species | Dilution | Company |
| --- | --- | --- | --- |
| **FLG** | Mouse | 1:100 | Thermo Fisher |
| **IVL** | Mouse | 1:20 | Mon 150 (own antibody) |
| **CLDN1** | Rabbit | 1:200 | Invitrogen |
| **CLDN4** | Mouse | 1:200 | Invitrogen |
| **Ki67** | Rabbit | 1:200 | Abcam |
| **SKALP** | Rabbit | 1:4000 | Own antibody |
| **KRT16** | Mouse | 1:200 | Santa-Cruz |
| **Goat anti-rabbit** | Goat | 1:200 | Vector laboratories |
| **Horse anti-mouse** | Mouse | 1:200 | Vector laboratories |

**Supplemental Figure 1:**
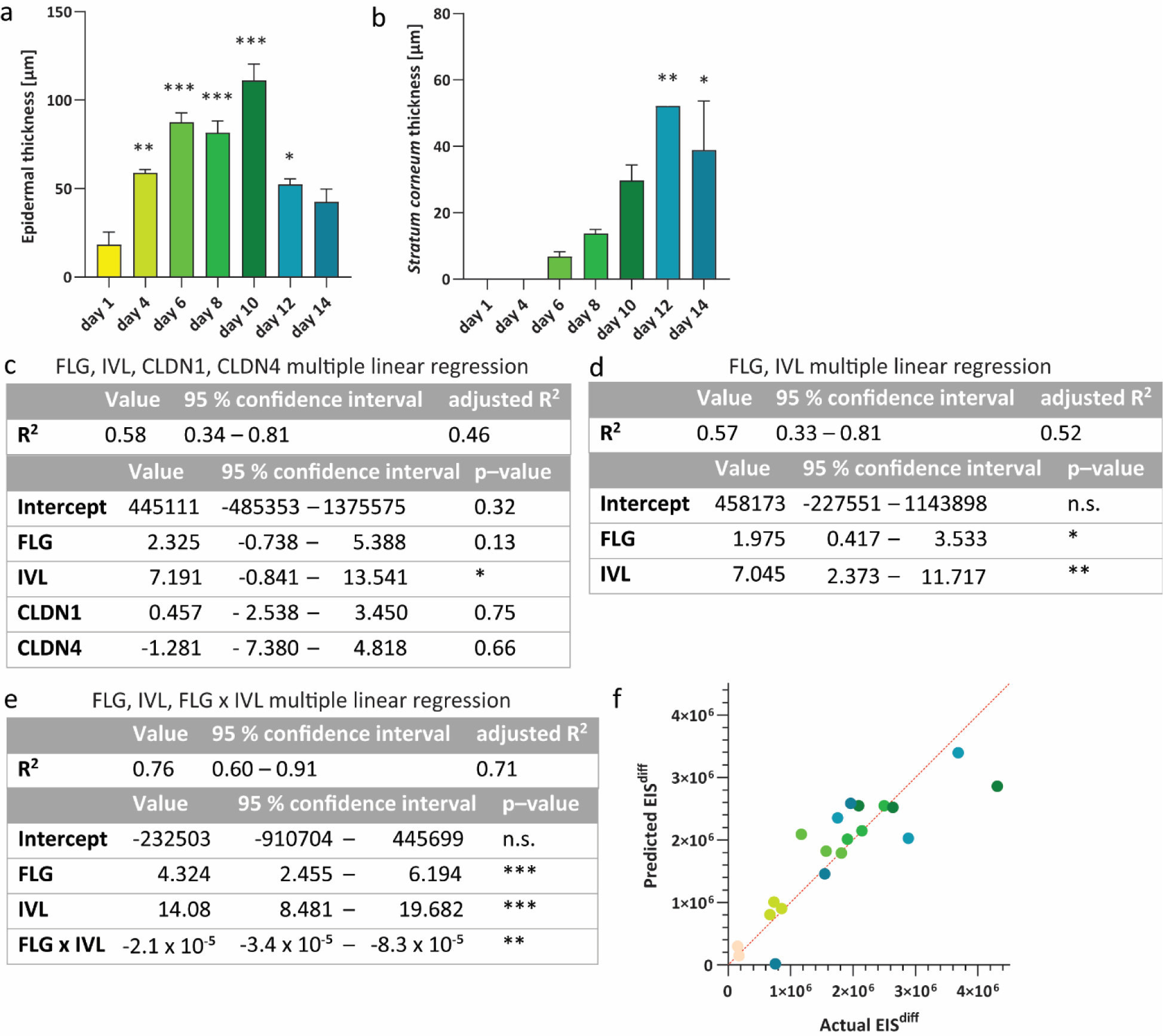
FLG, IVL and FLG x IVL interaction significantly influence EIS^diff^ during HEE development. (a‒b) (a) Epidermal thickness and (b) *stratum corneum* thickness of during HEE development compared to day 1. Each condition represents three biological replicates. (c‒e) Multiple linear regression using (c) FLG, IVL, CLDN1 and CLDN4, (d) FLG and IVL and (e) FLG, IVL and FLG x IVL expression to predict EIS^diff^. Analysis is based on all timepoints and replicates from figure 3. (f) Correlation of actual and FLG, IVL, FLG x IVL regressed EIS^diff^.

**Supplemental Figure 2:**
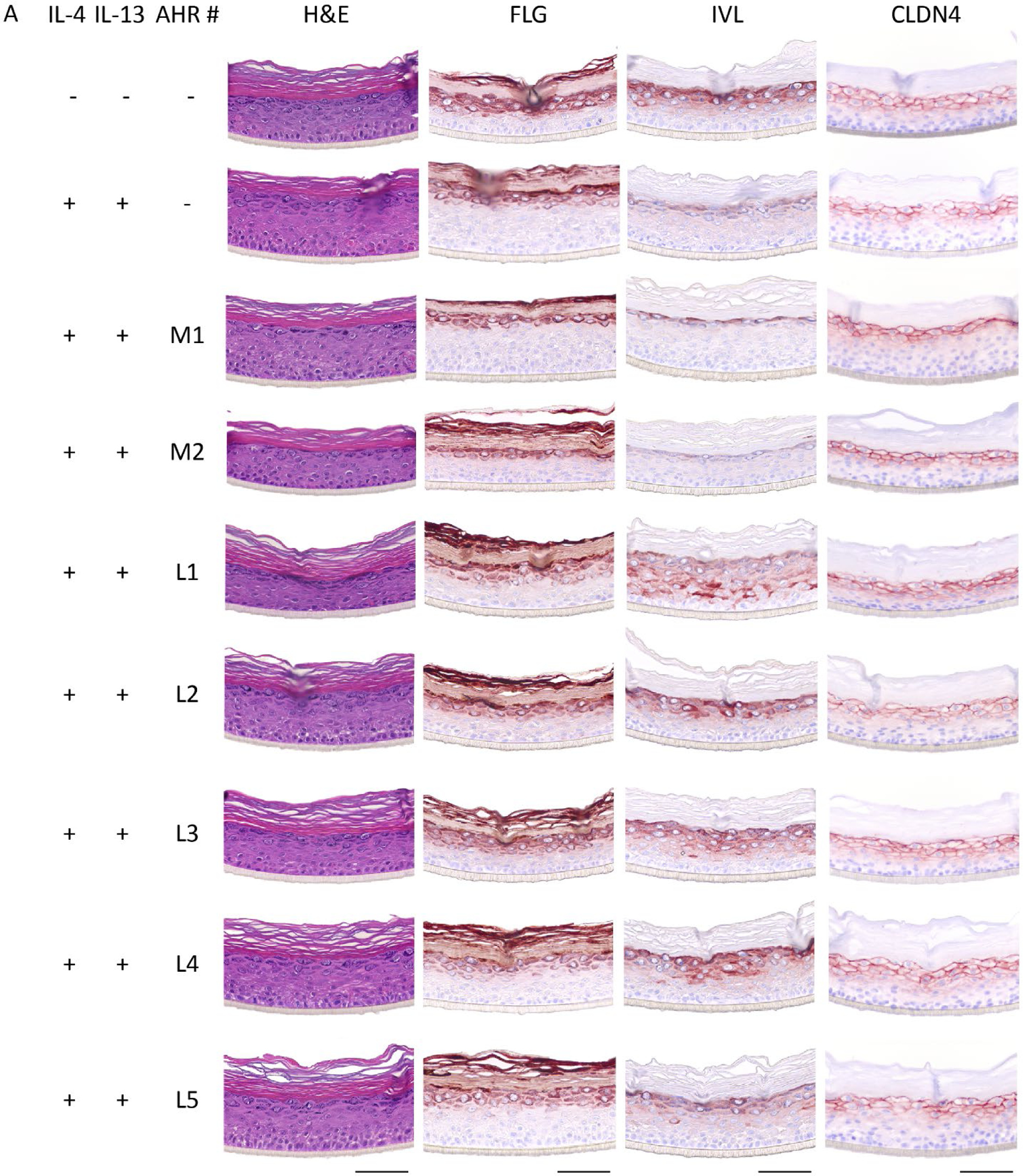
AHR activation mediates therapeutic response in a pro– inflammatory epidermis model. (a) HEEs stained for differentiation (FLG, IVL) and cell–cell adhesion (CLDN4) proteins. Pictures represent three biological replicates of HEEs at day 8 of air exposure and were taken with 40x magnification. Size bars indicate 100 µm.

## REFERENCES

Alexander H, Brown S, Danby S, Flohr C. Research Techniques Made Simple: Transepidermal Water Loss Measurement as a Research Tool. J. Invest. Dermatol. Elsevier; 2018. p. 2295–2300.e1

Arlk YB, Van Der Helm MW, Odijk M, Segerink LI, Passier R, Van Den Berg A, et al. Barriers-on-chips: Measurement of barrier function of tissues in organs-on-chips. Biomicrofluidics. American Institute of Physics; 2018;12(4)

Bäsler K, Bergmann S, Heisig M, Naegel A, Zorn-Kruppa M, Brandner JM. The role of tight junctions in skin barrier function and dermal absorption. J. Control. Release. J Control Release; 2016;242:105–18

Benson K, Cramer S, Galla HJ. Impedance-based cell monitoring: Barrier properties and beyond [Internet]. Fluids Barriers CNS. BioMed Central; 2013 [cited 2020 Dec 22]. p. 5

Bissonnette R, Stein Gold L, Rubenstein DS, Tallman AM, Armstrong A. Tapinarof in the treatment of psoriasis: A review of the unique mechanism of action of a novel therapeutic aryl hydrocarbon receptor–modulating agent. J. Am. Acad. Dermatol. Mosby Inc.; 2021. p. 1059–67

Van Den Bogaard EH, Bergboer JGM, Vonk-Bergers M, Van Vlijmen-Willems IMJJ, Hato S V., Van Der Valk PGM, et al. Coal tar induces AHR-dependent skin barrier repair in atopic dermatitis. J. Clin. Invest. American Society for Clinical Investigation; 2013;123(2):917–27

van den Bogaard EHJ, van Geel M, van Vlijmen-Willems IMJJ, Jansen PAM, Peppelman M, van Erp PEJ, et al. Deficiency of the human cysteine protease inhibitor cystatin M/E causes hypotrichosis and dry skin. Genet. Med. Nature Publishing Group; 2019;21(7):1559–67

van den Bogaard E, Ilic D, Dubrac S, Tomic-Canic M, Bouwstra J, Celli A, et al. Perspective and Consensus Opinion: Good Practices for Using Organotypic Skin and Epidermal Equivalents in Experimental Dermatology Research. J. Invest. Dermatol. Elsevier B.V.; 2021;141(1):203–5

Breternitz M, Kowatzki D, Langenauer M, Elsner P, Fluhr JW. Placebo-controlled, double-blind, randomized, prospective study of a glycerol-based emollient on eczematous skin in atopic dermatitis: Biophysical and clinical evaluation. Skin Pharmacol. Physiol. S. Karger AG; 2008;21(1):39–45

Brewer MG, Yoshida T, Kuo FI, Fridy S, Beck LA, De Benedetto A. Antagonistic effects of IL-4 on IL-17A-mediated enhancement of epidermal tight junction function. Int. J. Mol. Sci. MDPI AG; 2019;20(17)

Celebi Sözener Z, Cevhertas L, Nadeau K, Akdis M, Akdis CA. Environmental factors in epithelial barrier dysfunction. J. Allergy Clin. Immunol. Mosby; 2020;145(6):1517–28

Chamlin SL, Kao J, Frieden IJ, Sheu MY, Fowler AJ, Fluhr JW, et al. Ceramide-dominant barrier repair lipids alleviate childhood atopic dermatitis: Changes in barrier function provide a sensitive indicator of disease activity. J. Am. Acad. Dermatol. Mosby Inc.; 2002;47(2):198–208

Chen S, Einspanier R, Schoen J. Transepithelial electrical resistance (TEER): a functional parameter to monitor the quality of oviduct epithelial cells cultured on filter supports. Histochem. Cell Biol. Springer; 2015;144(5):509–15

Crowe A, Yue W. Semi-quantitative Determination of Protein Expression Using Immunohistochemistry Staining and Analysis: An Integrated Protocol. BIO-PROTOCOL. Bio-protocol, LLC; 2019;9(24)

Dickson MA, Hahn WC, Ino Y, Ronfard V, Wu JY, Weinberg RA, et al. Human keratinocytes that express hTERT and also bypass a p16(INK4a)-enforced mechanism that limits life span become immortal yet retain normal growth and differentiation characteristics. Mol. Cell. Biol. Mol Cell Biol; 2000;20(4):1436–47

Évora AS, Adams MJ, Johnson SA, Zhang Z. Corneocytes: Relationship between structural and biomechanical properties [Internet]. Skin Pharmacol. Physiol. S. Karger AG; 2021. p. 146–61

Eyerich S, Eyerich K, Traidl-Hoffmann C, Biedermann T. Cutaneous Barriers and Skin Immunity: Differentiating A Connected Network. Trends Immunol. Elsevier Current Trends; 2018. p. 315– 27

Faway E, Staerck C, Danzelle C, Vroomen S, Courtain C, Mignon B, et al. Towards a standardized procedure for the production of infective spores to study the pathogenesis of dermatophytosis. J. Fungi. MDPI; 2021;7(12):1029

Fernandes J, Karra N, Bowring J, Reale R, James J, Blume C, et al. Real-time monitoring of epithelial barrier function by impedance spectroscopy in a microfluidic platform. Lab Chip. Royal Society of Chemistry; 2022;22(10):2041–54

Fletcher TD. psychometric: Applied Psychometric Theory. R Packag. version 2.4. 2023.

Flohr C, England K, Radulovic S, McLean WHI, Campbell LE, Barker J, et al. Filaggrin loss-of-function mutations are associated with early-onset eczema, eczema severity and transepidermal water loss at 3 months of age. Br. J. Dermatol. Br J Dermatol; 2010;163(6):1333–6

Furue M. Regulation of filaggrin, loricrin, and involucrin by IL-4, IL-13, IL-17A, IL-22, AHR, and NRF2: Pathogenic implications in atopic dermatitis. Int. J. Mol. Sci. Multidisciplinary Digital Publishing Institute; 2020. p. 1–25

El Ghalbzouri A, Siamari R, Willemze R, Ponec M. Leiden reconstructed human epidermal model as a tool for the evaluation of the skin corrosion and irritation potential according to the ECVAM guidelines. Toxicol. Vitr. Pergamon; 2008;22(5):1311–20

Groeber F, Engelhardt L, Egger S, Werthmann H, Monaghan M, Walles H, et al. Impedance Spectroscopy for the Non-Destructive Evaluation of in Vitro Epidermal Models. Pharm. Res. Springer New York LLC; 2015;32(5):1845–54

Hadj-Rabia S, Baala L, Vabres P, Hamel-Teillac D, Jacquemin E, Fabre M, et al. Claudin-1 gene mutations in neonatal sclerosing cholangitis associated with ichthyosis: A tight junction disease. Gastroenterology. W.B. Saunders; 2004;127(5):1386–90

Haorah J, Schall K, Ramirez SH, Persidsky Y. Activation of protein tyrosine kinases and matrix metalloproteinases causes blood-brain barrier injury: Novel mechanism for neurodegeneration associated with alcohol abuse. Glia. NIH Public Access; 2008;56(1):78–88

Al Kindi A, Williams H, Matsuda K, Alkahtani AM, Saville C, Bennett H, et al. Staphylococcus aureus second immunoglobulin-binding protein drives atopic dermatitis via IL-33. J. Allergy Clin. Immunol. Mosby; 2021;147(4):1354–1368.e3

Kirschner N, Rosenthal R, Günzel D, Moll I, Brandner JM. Tight junctions and differentiation - a chicken or the egg question? Exp. Dermatol. Exp Dermatol; 2012;21(3):171–5

Klotz T, Ibrahim A, Maddern G, Caplash Y, Wagstaff M. Devices measuring transepidermal water loss: A systematic review of measurement properties. Ski. Res. Technol. John Wiley & Sons, Ltd; 2022;28(4):497–539

Liu X, Michael S, Bharti K, Ferrer M, Song MJ. A biofabricated vascularized skin model of atopic dermatitis for preclinical studies. Biofabrication. Institute of Physics Publishing; 2020;12(3):035002

Luengo J, Schneider M, Schneider AM, Lehr CM, Schaefer UF. Human skin permeation enhancement using plga nanoparticles is mediated by local ph changes. Pharmaceutics. Multidisciplinary Digital Publishing Institute; 2021;13(10):1608

Magar HS, Hassan RYA, Mulchandani A. Electrochemical impedance spectroscopy (Eis): Principles, construction, and biosensing applications. Sensors. Multidisciplinary Digital Publishing Institute (MDPI); 2021.

Mannweiler R, Bergmann S, Vidal-y-Sy S, Brandner JM, Günzel D. Direct assessment of individual skin barrier components by electrical impedance spectroscopy. Allergy Eur. J. Allergy Clin. Immunol. John Wiley and Sons Inc; 2021;76(10):3094–106

Morin M, Ruzgas T, Svedenhag P, Anderson CD, Ollmar S, Engblom J, et al. Skin hydration dynamics investigated by electrical impedance techniques in vivo and in vitro. Sci. Reports 2020 101. Nature Publishing Group; 2020;10(1):1–18

Natsuga K. Epidermal barriers. Cold Spring Harb. Perspect. Med. Cold Spring Harbor Laboratory Press; 2014 [cited 2021 May 26].

Nemoto-Hasebe I, Akiyama M, Nomura T, Sandilands A, McLean WHI, Shimizu H. Clinical severity correlates with impaired barrier in filaggrin-related eczema. J. Invest. Dermatol. Elsevier; 2009;129(3):682–9

Neupane R, Boddu SHS, Renukuntla J, Babu RJ, Tiwari AK. Alternatives to biological skin in permeation studies: Current trends and possibilities. Pharmaceutics. 2020.

Niehues H, Bouwstra JA, El Ghalbzouri A, Brandner JM, Zeeuwen PLJM, van den Bogaard EH. 3D skin models for 3R research: The potential of 3D reconstructed skin models to study skin barrier function. Exp. Dermatol. Blackwell Publishing Ltd; 2018. p. 501–11

Niehues H, Rikken G, van Vlijmen-Willems IMJJ, Rodijk-Olthuis D, van Erp PEJ, Zeeuwen PLJM, et al. Identification of Keratinocyte Mitogens: Implications for Hyperproliferation in Psoriasis and Atopic Dermatitis. JID Innov. Ski. Sci. from Mol. to Popul. Heal. JID Innov; 2021;2(1):100066

O’Donnell EF, Saili KS, Koch DC, Kopparapu PR, Farrer D, Bisson WH, et al. The anti-inflammatory drug leflunomide is an agonist of the aryl hydrocarbon receptor. PLoS One. Public Library of Science; 2010;5(10):e13128

Orsmond A, Bereza-Malcolm L, Lynch T, March L, Xue M. Skin barrier dysregulation in psoriasis. Int. J. Mol. Sci. Multidisciplinary Digital Publishing Institute (MDPI); 2021.

Pecoraro B, Tutone M, Hoffman E, Hutter V, Almerico AM, Traynor M. Predicting Skin Permeability by Means of Computational Approaches: Reliability and Caveats in Pharmaceutical Studies. J. Chem. Inf. Model. American Chemical Society; 2019. p. 1759–71

Poland A, Glover E. Stereospecific, high affinity binding of 2,3,7,8-tetrachlorodibenzo-p-dioxin by hepatic cytosol. J. Biol. Chem. 1976;251(16):4936–46

R Core Team (2023). R: A language and environment for statistical computing. R foundation for statistical computing. https://www.R-project.org/. 2023.

Riethmüller C. Assessing the skin barrier via corneocyte morphometry [Internet]. Exp. Dermatol. John Wiley & Sons, Ltd; 2018. p. 923–30

Rikken G, van den Brink NJM, van Vlijmen-Willems IMJJ, van Erp PEJ, Pettersson L, Smits JPH, et al. Carboxamide Derivatives Are Potential Therapeutic AHR Ligands for Restoring IL-4 Mediated Repression of Epidermal Differentiation Proteins. Int. J. Mol. Sci. MDPI; 2022;23(3)

Rikken G, Smith KJ, van den Brink NJM, Smits JPH, Gowda K, Alnemri A, et al. Lead optimization of aryl hydrocarbon receptor ligands for treatment of inflammatory skin disorders. Biochem. Pharmacol. Biochem Pharmacol; 2023;208:115400

Rinaldi AO, Korsfeldt A, Ward S, Burla D, Dreher A, Gautschi M, et al. Electrical impedance spectroscopy for the characterization of skin barrier in atopic dermatitis. Allergy Eur. J. Allergy Clin. Immunol. John Wiley & Sons, Ltd; 2021;76(10):3066–79

Roberts MS, Cheruvu HS, Mangion SE, Alinaghi A, Benson HAE, Mohammed Y, et al. Topical drug delivery: History, percutaneous absorption, and product development. Adv. Drug Deliv. Rev. Adv Drug Deliv Rev; 2021.

Safe SH. Comparative Toxicology and Mechanism of Action of Polychlorinated Dibenzo-P-Dioxins and Dibenzofurans. Annu. Rev. Pharmacol. Toxicol. Annual Reviews 4139 El Camino Way, P.O. Box 10139, Palo Alto, CA 94303-0139, USA; 1986;26(1):371–99

Shamaprasad P, Frame CO, Moore TC, Yang A, Iacovella CR, Bouwstra JA, et al. Using molecular simulation to understand the skin barrier. Prog. Lipid Res. Pergamon; 2022. p. 101184

van Smeden J, Janssens M, Gooris GS, Bouwstra JA. The important role of stratum corneum lipids for the cutaneous barrier function. Biochim. Biophys. Acta - Mol. Cell Biol. Lipids. Elsevier; 2014;1841(3):295–313

Smits JPH, van den Brink NJM, Meesters LD, Hamdaoui H, Niehues H, Jansen PAM, et al. Investigations into the filaggrin null phenotype: showcasing the methodology for CRISPR/Cas9 editing of human keratinocytes. J. Invest. Dermatol. J Invest Dermatol; 2023a

Smits JPH, Niehues H, Rikken G, Van Vlijmen-Willems IMJJ, Van De Zande GWHJF, Zeeuwen PLJM, et al. Immortalized N/TERT keratinocytes as an alternative cell source in 3D human epidermal models. Sci. Rep. 2017;7(1):1–14

Smits JPH, Qu J, Pardow F, van den Brink NJM, Rodijk-Olthuis D, van Vlijmen-Willems IMJJ, et al. The aryl hydrocarbon receptor regulates epidermal differentiation through transient activation of TFAP2A. bioRxiv Prepr. Serv. Biol. bioRxiv; 2023b

Srinivasan B, Kolli AR, Esch MB, Abaci HE, Shuler ML, Hickman JJ. TEER Measurement Techniques for In Vitro Barrier Model Systems. J. Lab. Autom. NIH Public Access; 2015. p. 107– 26

Sugarman JL, Fluhr JW, Fowler AJ, Bruckner T, Diepgen TL, Williams ML. The Objective Severity Assessment of Atopic Dermatitis Score: An Objective Measure Using Permeability Barrier Function and Stratum Corneum Hydration with Computer-Assisted Estimates for Extent of Disease. Arch. Dermatol. Arch Dermatol; 2003;139(11):1417–22

Sugawara T, Iwamoto N, Akashi M, Kojima T, Hisatsune J, Sugai M, et al. Tight junction dysfunction in the stratum granulosum leads to aberrant stratum corneum barrier function in claudin-1-deficient mice. J. Dermatol. Sci. J Dermatol Sci; 2013;70(1):12–8

Tárnoki-Zách J, Mehes E, Varga-Medveczky Z, Isai DG, Barany N, Bugyik E, et al. Development and evaluation of a human skin equivalent in a semiautomatic microfluidic diffusion chamber. Pharmaceutics. Multidisciplinary Digital Publishing Institute (MDPI); 2021;13(6)

Tjabringa G, Bergers M, Van Rens D, De Boer R, Lamme E, Schalkwijk J. Development and validation of human psoriatic skin equivalents. Am. J. Pathol. Elsevier; 2008;173(3):815–23

Wegener J, Hakvoort A, Galla HJ. Barrier function of porcine choroid plexus epithelial cells is modulated by cAMP-dependent pathways in vitro. Brain Res. Elsevier; 2000;853(1):115–24

Yang G, Seok JK, Kang HC, Cho YY, Lee HS, Lee JY. Skin barrier abnormalities and immune dysfunction in atopic dermatitis. Int. J. Mol. Sci. Multidisciplinary Digital Publishing Institute; 2020. p. 2867

Yeste J, Illa X, Alvarez M, Villa R. Engineering and monitoring cellular barrier models. J. Biol. Eng. BioMed Central; 2018.

Yoshida T, Beck LA, De Benedetto A. Skin barrier defects in atopic dermatitis: From old idea to new opportunity. Allergol. Int. Elsevier; 2022. p. 3–13

